# Correcting model misspecification in relationship estimates

**DOI:** 10.1101/2024.05.13.594005

**Authors:** Ethan M. Jewett, the 23andMe Research Team

## Abstract

The datasets of large genotyping biobanks and direct-to-consumer genetic testing companies contain many related individuals. Until now, it has been widely accepted that the most distant relationships that can be detected are around fifteen degrees (approximately 8^*th*^ cousins) and that practical relationship estimates have a ceiling around ten degrees (approximately 5^*th*^ cousins). However, we show that these assumptions are incorrect and that they are due to a misapplication of relationship estimators. In particular, relationship estimators are applied almost exclusively to putative relatives who have been identified because they share detectable tracts of DNA identically by descent (IBD). However, no existing relationship estimator conditions on the event that two individuals share at least one detectable segment of IBD anywhere in the genome. As a result, the relationship estimates obtained using existing estimators are dramatically biased for distant relationships, inferring all sufficiently distant relationships to be around ten degrees regardless of the depth of the true relationship. Existing relationship estimators are derived under a model that assumes that each pair of related individuals shares a single common ancestor (or mating pair of ancestors). This model breaks down for relationships beyond 10 generations in the past because individuals share many thousands of cryptic common ancestors due to pedigree collapse. We first derive a corrected likelihood that conditions on the event that at least one segment is observed between a pair of putative relatives and we demonstrate that the corrected likelihood largely eliminates the bias in estimates of pairwise relationships and provides a more accurate characterization of the uncertainty in these estimates. We then reformulate the relationship inference problem to account for the fact that individuals share many common ancestors, not just one. We demonstrate that the most distant relationship that can be inferred using IBD may be 200 degrees or more, rather than ten, extending the time-to-common ancestor from approximately 300 years in the past to approximately 3,000 years in the past or more. This dramatic increase in the range of relationship estimators makes it possible to infer relationships whose common ancestors lived before historical events such as European settlement of the Americas, the Transatlantic Slave Trade, and the rise and fall of the Roman Empire.

## 2. Introduction

A genetic relationship inference method is an algorithm that takes as input the genotyped or sequenced DNA of a putative pair of relatives and potentially other information such as ages and sexes and returns an estimate of their relationship. These algorithms are commonly applied in the context of direct-to-consumer genetic testing in order to identify relatives and infer genealogies [Henn et al., 2012, Ball et al., 2016, Jewett et al., 2021] and they are applied in the context of medical genetic studies to identify and leverage or control for cryptic relatedness [Voight and Pritchard, 2005, Staples et al., 2018, Howe et al., 2022].

All relationship inference methods rely on probability distributions that describe how much IBD is observed between a pair of individuals as a function of their relationship. Common statistics include the total length of observed IBD in centimorgans [Henn et al., 2012, Ball et al., 2016, Jewett et al., 2021] or equivalent quantities such as the kinship coefficient [Staples et al., 2014, 2016, Manichaikul et al., 2010, Ramstetter et al., 2018]. Other common quantities include the number of observed IBD segments and their lengths [Huff et al., 2011].

Regardless of the approach, all existing methods rely implicitly or explicitly on distributions of IBD statistics that were obtained without conditioning on the event that IBD was observed between the two putative relatives. Likelihood methods like the ERSA method of Huff et al. [2011] rely on the probability distribution of one or more observed IBD statistics. Estimators commonly used in direct-to-consumer (DTC) genetic testing rely on empirical versions of these probability distributions that can be obtained using simulations [Henn et al., 2012, Ball et al., 2016]. Other methods rely on analytically-derived bounds that delineate regions of “IBD space” in which the observed values of IBD statistics are most consistent with different relationships [Manichaikul et al., 2010, Ramstetter et al., 2018].

All of these distributions are unconditional on IBD sharing. Although Huff et al. [2011] derive a conditional version of the probability distribution of observed segment lengths conditional on the event that two putative relatives are ascertained because they share IBD at a particular locus, this distribution is not the same as the distribution conditional on observing at least one segment of IBD anywhere in the genome.

Estimates made without conditioning on observing at least one IBD segment are appropriate in scenarios in which pairs of putative relatives were ascertained in a manner that does not depend on the amount of IBD they share; for example, to verify a self-reported relationship that was identified based on previous genealogical knowledge. However, in most contexts in which relationship inference is applied, pairs of putative relatives are ascertained by first detecting shared IBD. In this context, it is inappropriate to apply estimators that do not condition on the event that IBD is observed.

Failure to condition on the observation of IBD has relatively little effect on close relationships because the probability that close relatives share IBD is high. However, when relationships are distant, failure to condition on the event that IBD is observed has profound consequences resulting in dramatically biased relationship estimates as we will demonstrate.

Here, we derive the probability distribution of the observed number of IBD segments and their lengths as a function of the genealogical relationship that gave rise to the segments, conditional on the event that at least one segment of IBD was observed. We show that the corrected estimator no longer has the profound bias observed in the unconditional estimator and that it allows relationship estimates that extend into the distant past.

We also derive a relationship estimator that explicitly accounts for the fact that pairs of individuals have many thousands of common ancestors. This model of relatedness is arguably more realistic than the most prevalent existing model of relatedness in which each pair of individuals has exactly one common ancestor or mating pair of common ancestors.

Finally, we derive a approximate formula for the expected number of ancestors in each generation in the past who contributed a detectable IBD segment longer than a minimum segment length to a given pair of present-day individuals. Using this formula, we demonstrate that the number of detectable-IBD-contributing common ancestors living more than 100 generations (∼3,000 years) in the past is likely to be non-negligible.

## 3. Results

### 3.1. The expected fraction of relationships that are beyond the range of existing estimators

Before investigating the bias in existing relationship estimators, it’s informative to consider how often we might expect to encounter distant IBD-sharing relationships in the first place. If each each pair of relatives had exactly one common ancestor (or mating pair of common ancestors), then the probability of sharing IBD with a distant relative would indeed be very small. Caballero et al. [2019] found that simulated sixth cousins with just one pair of common ancestors typically shared no IBD segment with one another, and a related analysis found that simulated 8th cousins and beyond (individuals who shared a pair of common ancestors nine generations or more in the past) were exceedingly unlikely to share any detectable IBD segments at all [Williams, 2024]. For instance, the probability that 8th cousins with a single pair of common ancestors shared at least one segment was less than 1%. Henn et al. [2012] predicted that IBD from simulated 9th cousins and beyond would be undetectable (Table 2 of Henn et al. [2012]).

Although it is very unlikely for two distant relatives to share detectable IBD through a particular common ancestor, each pair of individuals has many common genealogical ancestors (Figure 1). Moreover, due to pedigree collapse, each modern individual can have multiple semi-independent lineages back to each of their common ancestors. Thus, the chances of observing a very distant IBD-sharing relationship may actually be quite high.

**Figure 1.**
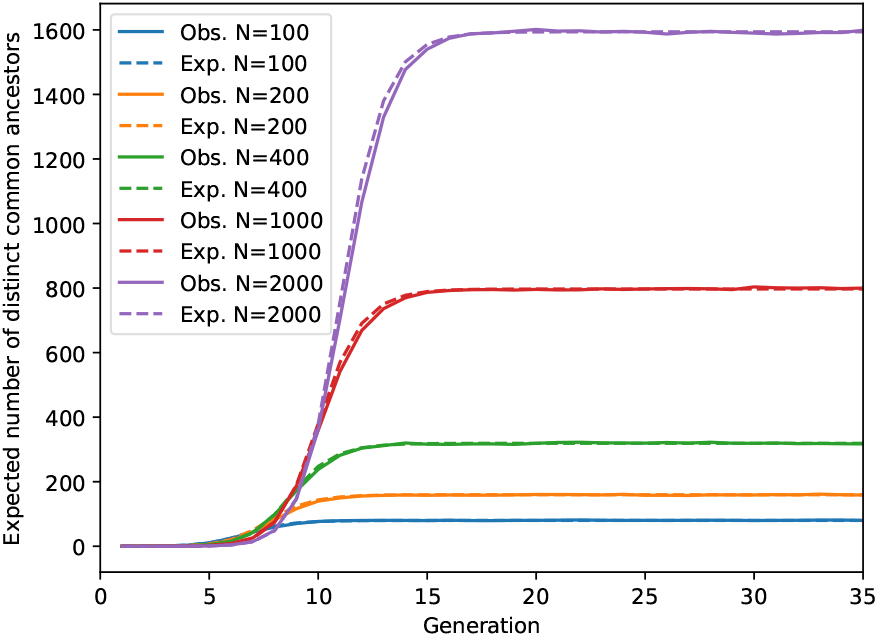
The expected number of distinct common ancestors shared between two present-day people. The simulated values are compared with analytical values obtained using Equation (29) for various population sizes for which simulation is fast.

Although a pair of present-day people may have many genealogical common ancestors, only some of these ancestors contributed detectable segments of IBD longer than the minimum observable segment length *τ* cM. Figure 2A shows the expected number of distinct detectable-IBD-transmitting common ancestors in generation *g* in the past that are shared between two individuals in the present day. The analytical expectation is compared with the mean number of common ancestors observed in simulations for several population sizes that are small enough to be computationally efficient. The approximation is relatively good except for very small and very large generation times.

**Figure 2.**
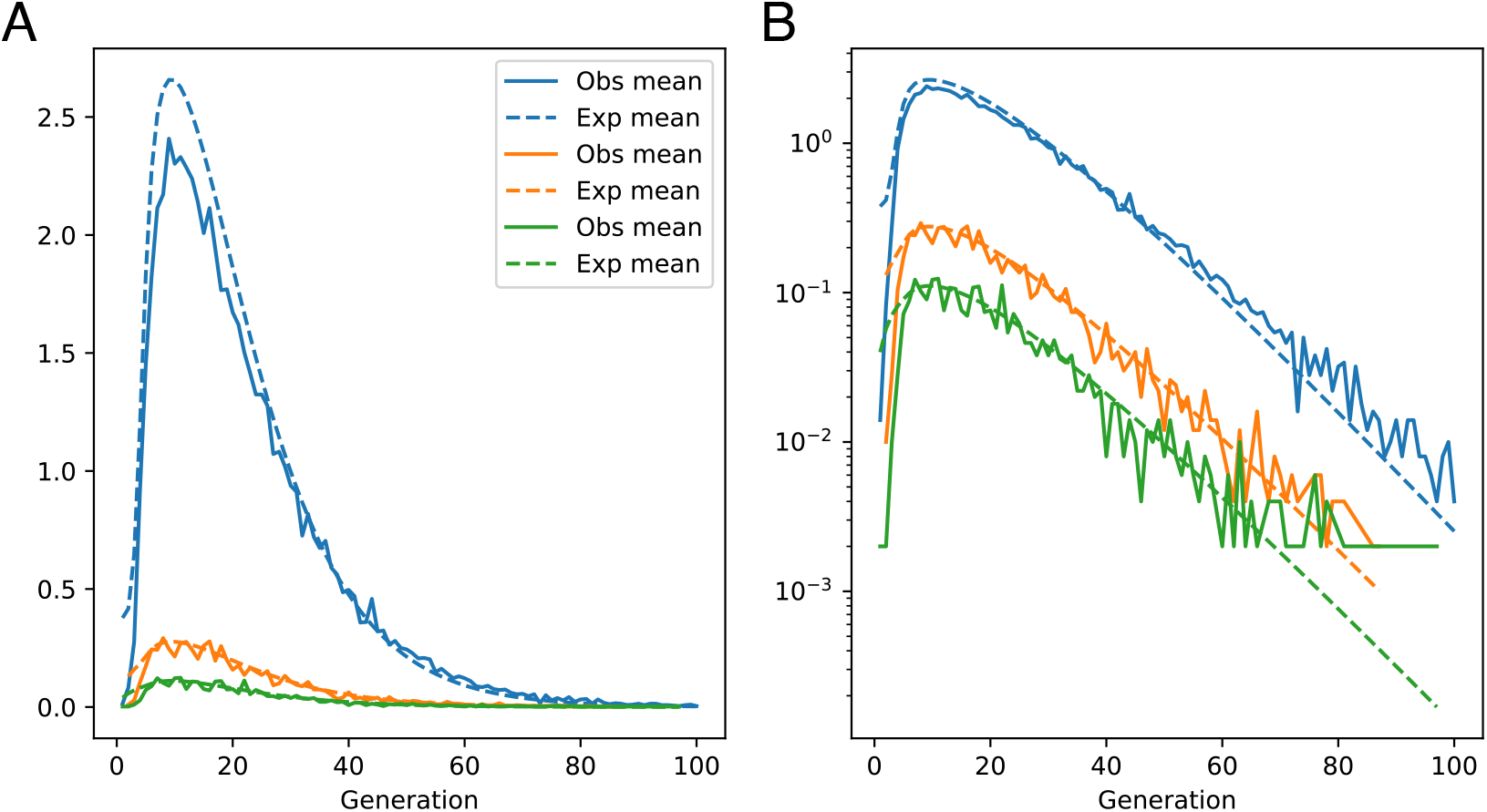
(A) The expected number of distinct detectable-IBD-transmitting common ancestors at each generation in the past. Curves are shown for a minimum segment length of *τ* = 5 cM and for three different effective population sizes: *N* = 1, 000, *N* = 2, 000 and *N* = 5, 000 individuals. (B) Same as (A) in log scale.

At small generation times, the analytical expectation is an overestimate because the coalescent is not restricted by the pedigree; it therefore allows segments to find common ancestors amongst the full population even one generation in the past. However the approximation becomes fairly accurate even a few generations in the past. One could correct for this discrepancy between the analytical and simulated values for small generation times; however, the discrepancy is unlikely to have much effect on relationship inference because the information for inferring close relatives is so strong that it overcomes the fact that the prior is slightly misspecified for close relationships (Figure 3D).

**Figure 3.**
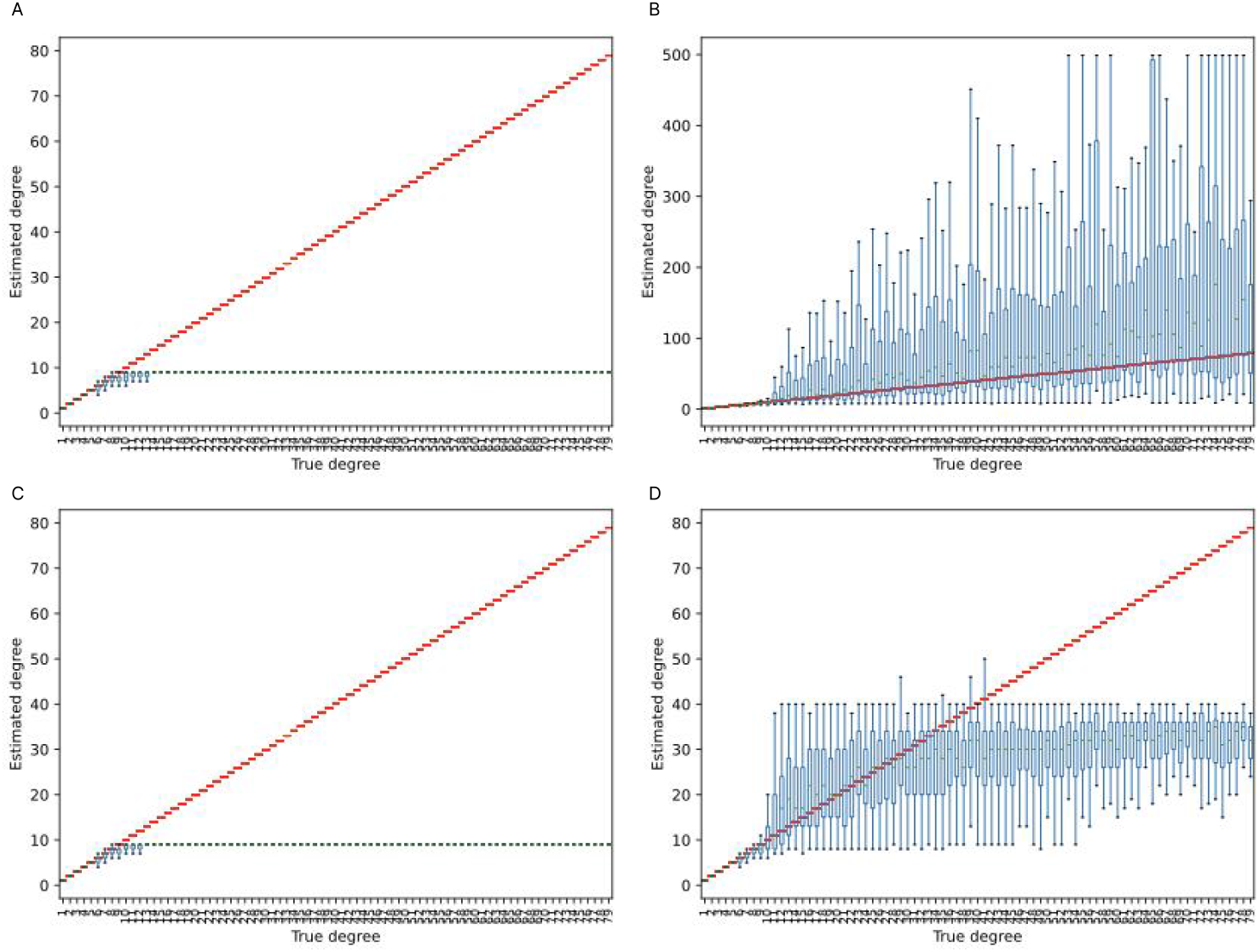
Inferred degree using unconditional and conditional estimators for relationships between 1 and 79 degrees. (A) The unconditional likelihood. (B) The conditional likelihood (Equation 8). (C) The unconditional likelihood together with the prior obtained by normalizing Equation (19) with *N* = 10, 000. (D) The conditional likelihood together with the prior obtained by normalizing Equation (19) with *N* = 10, 000.

At large generation times and for large population sizes, the simulations predict more shared ancestors than the analytical approximation, perhaps because there is a non-negligible probability of observing two adjacent segments. Such adjacent segments have a higher probability of being long enough to be observed. As in the case of small coalescent times, the discrepancy for large coalescent times is unlikely to have much effect on estimates because the peak of the prior occurs around 10 generations and forces most Bayesian estimates to be recent (Figure 3D).

From Figure 2, it can be seen that the probability of observing a detectable IBD segment longer than 5 cM from an ancestor who lived many generations in the past is non-negligible. For a particular pair of present-day individuals, Table 1 quantifies the expected number of detectable-IBD-contributing common ancestors living at least *g* generations in the past in a population with effective population size *N*. From Table 1, it can be seen that the expected number of ancestors arising at least 100 generations in the past is at least 10^−4^, even in populations with relatively large effective sizes.

**TABLE 1.** Expected number of detectable-IBD-contributing common ancestors between two particular present-day individuals. Expectations are shown for various values of *g* and effective population sizes *N*.

| N | Generation |  |  |  |  |  |
| --- | --- | --- | --- | --- | --- | --- |
|  | 1 | 10 | 20 | 50 | 75 | 100 |
| 1,000 | 14.4159 | 10.9097 | 5.9837 | 0.5863 | 0.0673 | 0.0071 |
| 5,000 | 2.9688 | 2.2043 | 1.2128 | 0.1202 | 0.0139 | 0.0015 |
| 10,000 | 1.4921 | 1.1035 | 0.6074 | 0.0603 | 0.0070 | 0.0007 |
| 50,000 | 0.3000 | 0.2209 | 0.1216 | 0.0121 | 0.0014 | 0.0001 |
| 100,000 | 0.1501 | 0.1105 | 0.0608 | 0.0060 | 0.0007 | 0.0001 |

**TABLE 2.** Bounds *L*_*d*_ on the regions in which the likelihood is maximized for degree *d* whenever *L* ∈ (*L*_*d*+1_, *L*_*d*_]. Values are shown for the case *a* = 1 and *τ* = 0.

| m | Unconditional | Conditional | m | Unconditional | Conditional |
| --- | --- | --- | --- | --- | --- |
| 1 | 2267.06 | 2267.06 | 24 | - | 3.92 |
| 2 | 1275.18 | 1275.18 | 25 | - | 3.77 |
| 3 | 743.12 | 743.12 | 26 | - | 3.64 |
| 4 | 357.57 | 357.57 | 27 | - | 3.51 |
| 5 | 172.78 | 172.82 | 28 | - | 3.39 |
| 6 | 82.83 | 83.44 | 29 | - | 3.28 |
| 7 | 38.59 | 41.65 | 30 | - | 3.17 |
| 8 | 16.28 | 23.52 | 31 | - | 3.08 |
| 9 | 3.39 | 15.72 | 32 | - | 2.99 |
| 10 | 0. | 11.98 | 33 | - | 2.90 |
| 11 | - | 9.91 | 34 | - | 2.82 |
| 12 | - | 8.62 | 35 | - | 2.74 |
| 13 | - | 7.72 | 36 | - | 2.67 |
| 14 | - | 7.06 | 37 | - | 2.60 |
| 15 | - | 6.53 | 38 | - | 2.53 |
| 16 | - | 6.10 | 39 | - | 2.47 |
| 17 | - | 5.74 | 40 | - | 2.41 |
| 18 | - | 5.42 | 41 | - | 2.35 |
| 19 | - | 5.13 | 42 | - | 2.30 |
| 20 | - | 4.88 | 43 | - | 2.25 |
| 21 | - | 4.65 | 44 | - | 2.20 |
| 22 | - | 4.45 | 45 | - | 2.15 |
| 23 | - | 4.26 | 46 | - | 2.11 |
| 24 | - | 4.08 | ... | ... | ... |

Although pairs of individuals are not independent, the values in Table 1 suggest that each person can have thousands of distant detectable-IBD-contributing common ancestors with others in a database containing millions of individuals. Moreover, since the analytical approximation underestimates the expected number of distant ancestors, the expected number of distant ancestors is probably even higher.

The take-home message of Figure 2 and Table 1 is that IBD-sharing relationships greater than ten degrees are likely to occur frequently. These relationships are all inferred as tenth-degree relationships by existing relationship estimators. Therefore, we must use relationship estimators that are capable of inferring these distant relationships.

### 3.2. The degree of bias in existing estimators

To understand the bias that results when unconditional relationship estimators are applied to relatives ascertained on the basis of IBD sharing, we simulated IBD between relatives of various degrees, conditional on the event that they shared at least one detectable segment of IBD with one another [Jewett, 2024]. This sampling scheme reflects what we would expect to see in most real data applications involving relative detection in direct-to-consumer genetic testing or biobank data.

We sampled IBD for 1,000 relative pairs for each degree of relationship between one and forty degrees. For each simulation replicate, we inferred the degree of the relationship between the pair of individuals by maximizing the unconditional likelihood [Huff et al., 2011] and separately by maximizing the conditional likelihood derived in Section 5.1.

From Figure 3A, it can be seen that the unconditional relationship estimates are fairly accurate for close relationships up to approximately ten degrees (approximately fourth cousins), but they begin to diverge sharply from the true degree for relationships beyond ten degrees. Moreover, as the degree of the true relationship increases, the estimated relationships become increasingly tightly grouped around ten degrees. The ceiling at ten degrees is a property of the unconditional likelihood, as we discuss in Section 3.4. Figure 3 shows estimates when all IBD segments are detectable. However, setting a threshold on the minimum detectable segment length has little effect on the estimates, given that IBD is observed in the first place (Figure S1).

In contrast to the unconditional estimates, the conditional estimates shown in Figure 3B have considerably less bias. For these estimates, the mean inferred degree tracks reasonably well with the true degree (red line). The trade-off for reduced bias is increased variance, which can be seen in the range of inferred values shown in the boxplot. This increased variance is due to the fact that there is typically only one segment shared between distant relatives and all information about the degree of the relationship is contained in the length of that segment. As noted in Caballero et al. [2019], segment lengths for distant relationships do not carry much information about the true relationship degree, as the segment length distributions for different degrees overlap considerably.

### 3.3. Bayesian relationship estimates

The approximate prior distribution for the generation in which an IBD-contributing ancestor lived (Equation 19 and Figures 2C and 2D) can be used to obtain a Bayesian estimate of the relationship. The accuracy of the Bayesian estimate using the unconditional estimator is shown in Figure 3C and the accuracy of the conditional Bayesian estimator is shown in Figure 3D for an effective population size of *N* = 10, 000.

From Figures 3C and 3D, it can be seen that the unconditional estimator continues to have a ceiling at ten or eleven degrees. The prior introduces considerable bias into the conditional estimator as well, with the counterpoint that the variability in the estimates is dramatically reduced and all estimates are constrained to lie within a range that is more likely for human populations.

Ultimately, the prior has a considerable effect on the estimates so it is important to be fairly confident that the prior captures the true range of possible degrees of relationship. As we have noted, the prior in Equation (19) underestimates the number of ancestors observed at very deep timescales; however, even without this bias it seems unlikely that the prior would admit deep estimates beyond 50 generations or so. Thus, there is potentially an argument for employing the likelihood estimator for certain applications because it allows for deep estimates when the true relationship is distant, whereas the Bayesian estimator does not. It may therefore provides a less biased picture of overall relationship.

### 3.4. The total length of IBD

Figure 3 shows the inferred relationship using the full set of observed segment lengths. However, it is also common for relationship estimators to use the total sum 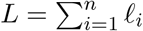 of lengths of observed IBD [Ball et al., 2016, Jewett et al., 2021] or a related statistic such as the kinship coefficient [Staples et al., 2014, 2016, Manichaikul et al., 2010, Ramstetter et al., 2018]. In Section 5.2, we derive a formula for the total observed length *L* of IBD for both the conditional case in which IBD was observed and the unconditional case in which IBD may or may not have been observed.

Figure 4A shows the distribution of the total length *L* of IBD for *a* = 1 common ancestors and several small values of *m*, the number of meioses in the lineage connecting two putative relatives. For small values of *m*, it can be seen that the unconditional and conditional distributions are nearly identical. This is because the probability of observing at least one segment of IBD is nearly one when *m* is small and the correction term 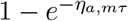 in Equation (14) is approximately one.

**Figure 4.**
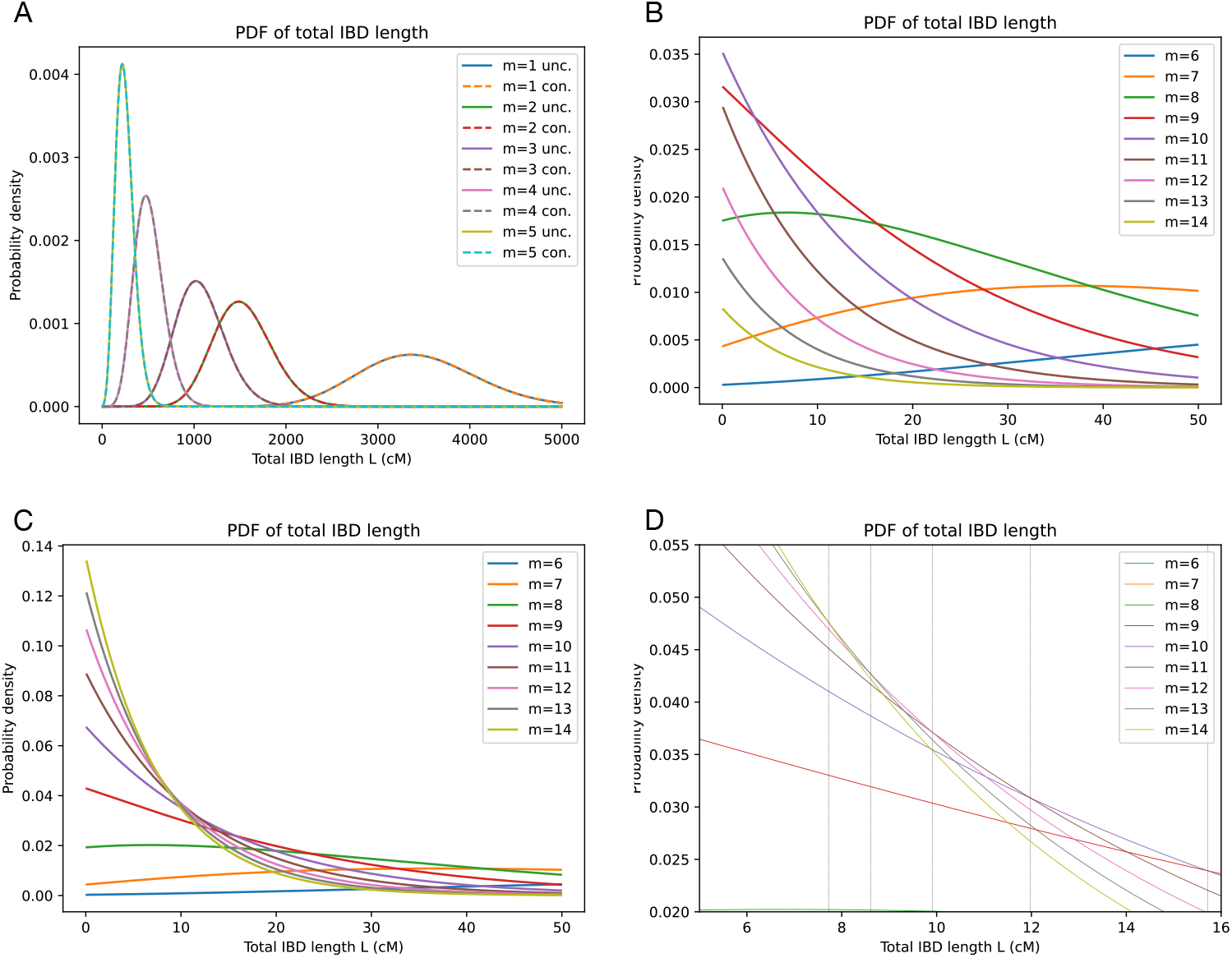
The distribution of the total length of IBD. (A) The distribution of the total length of IBD for *a* = 1 ancestors and several values of the number *m* of meioses separating two putative relatives. Both the unconditional (unc.) and conditional (con.) distributions are shown. (B) The unconditional distribution for values of *m* in the range *m* ∈ {6, …, 14}. (C) The conditional distribution for values of *m* in the range *m* ∈ {6, …, 14}. (D) Close up of the conditional distribution along with points *L*_*d*_ (black vertical lines) marking transition points where the likelihood surface for *d* = *m* − *a* + 1 is greater than the likelihood surface for *d* = *m* + 1 − *a* + 1.

The difference between the conditional and unconditional distributions can be seen by comparing Figures 4B and 4C. For values of *m* in the range 6 to 14, the conditional and unconditional distributions begin to diverge from one another and begin to take on qualitatively different behavior.

Figure 4B explains why the likelihood estimator in Figure 3A tops out at *d* = *m* − *a* + 1 ≈ 10 degrees. In particular, for *a* = 1 the density for all *m* > 10 is uniformly lower than the density for *m* = 10 in the region *L* > 0. This property of the density is made possible by the fact that the unconditional distribution has a point mass at *L* = 0, allowing the density for *L* > 0 to integrate to less than one. This property implies that the greatest possible value of *m* that can be inferred by maximum likelihood is *m* = 10 when *a* = 1 and *m* = 11 when *a* = 2 since the likelihood surface for all higher values of *m* is uniformly lower. Thus, the asymptote in Figures 3A and 3C at *d* = *m* − *a* + 1 = 10 degrees is a fundamental property of the likelihood. Moreover, all existing estimators do something similar to maximizing the unconditional likelihood, which results in a ceiling at *d* = 10 degrees for existing estimators.

In contrast to the unconditional likelihood, an estimator based on the conditional likelihood (Figures 4C and 4D) does not have a ceiling. The reason is that for any degree *d* > 0, there is always a region (*L*_*d*+1_, *L*_*d*_] such that the likelihood is maximized at degree *d* whenever the total sum of IBD lengths *L* is within (*L*_*d*+1_, *L*_*d*_]. The bounds *L*_*d*_ of these regions (black vertical lines) are shown in the close-up of the conditional distribution shown in Figure 4D.

### 3.5. Regions where the likelihood is maximized

A relationship estimator can search for the values of *a* and *m* that maximize the likelihood of the observed value of *L*; however, for a particular value of *a* it is also possible to precompute regions (*L*_*a,m*+1_, *L*_*a,m*_] such that the likelihood of *L* is maximized by *a* and *m* for *L* ∈ (*L*_*a,m*+1_, *L*_*a,m*_]. This is the approach taken by some genetic testing companies, where the regions (*L*_*a,m*+1_, *L*_*a,m*_] are determined empirically using simulations [Henn et al., 2012, Ball et al., 2016]. Other methods use similar bounds obtained from kinship coefficients [Manichaikul et al., 2010, Ramstetter et al., 2018]. In general, because there is more information for inferring the compound parameter *d* = *m* − *a* + 1 and it is difficult to resolve *a* and *m* for the same value of *d*, estimators typically express ranges in terms of *d* rather than *a* and *m*. Specifically, they use the ranges (*L*_*d*+1_, *L*_*d*_], which can be approximated by setting *a* to a fixed value.

Note that because the simulations that are used to obtain the regions (*L*_*d*+1_, *L*_*d*_] are performed unconditionally on the event *O* that an IBD segment is observed, the resulting estimator is equivalent to the maximum likelihood estimator based on the unconditional distribution (Figure 3A). Kinship coefficients are also unconditional on IBD sharing. In Section 5.3, we use the conditional distribution of *L* to derive the bounds (*L*_*d*+1_, *L*_*d*_] under the conditional likelihood. These are shown in Table 2.

From Table 2, we can see that *d* = *m* = 10 is the most likely degree in the region *L* ∈ [0, 3.39] for the unconditional estimator and the full region *L* > 0 is covered by regions corresponding to degrees 1 through 10. In comparison, for the conditional likelihood, there is a region of *L*-space in which each degree *d* is the most likely degree.

## 4. A relationship estimator that accounts for multiple ancestors

So far, we have considered the problem of updating existing relationship estimators to condition on the event that at least one segment of IBD is observed. However, these estimators make use of a conceptual model of relatedness in which each pair of individuals is connected through a single common ancestor or mating pair of common ancestors (5A). Individuals *i* and *j* in Figure 5A may have other very distant ancestors (grey circles) that give rise to occasional small segments of “background IBD,” but in this conceptual model, “background IBD” reflects very distant relationships that we aren’t interested in.

**Figure 5.**
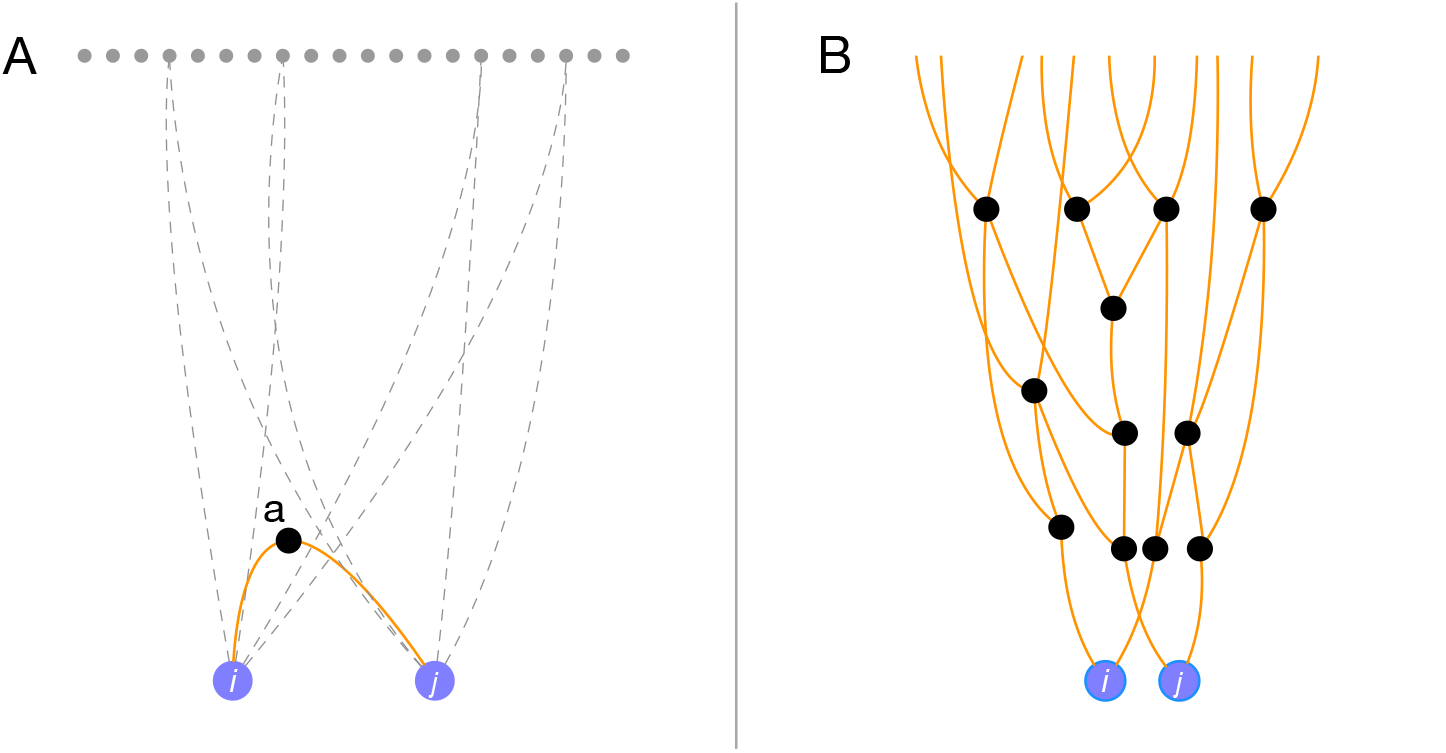
Conceptual models for the development of relationship estimators. Panel A shows the conceptual model that underlies existing relationship estimators. In this model, each pair of individuals, *i* and *j* (purple dots), shares a single common ancestor, *a*, or a single mating pair of common ancestors (*a*_1_, *a*_2_) (black circle). Other genealogical ancestors (grey dots) exist in the very distant past, but any IBD these ancestors contribute amounts to background noise. Panel B shows a conceptual model that more accurately describes genealogical relatedness at distant timescales. In this model, each pair of individuals shares many common ancestors in each generation in the past. Some of these ancestors contribute detectable IBD to the pair and some do not.

The assumption that two people are connected through a single close relationship may be true when we restrict our attention to the very recent past (perhaps within the most recent five to ten generations), but this assumption quickly breaks down as the degree of the relationship increases. As we discussed in Section 3.1, the number of common genealogical ancestors between two people can be large even in the not-too-distant past and each individual has many lineages connecting them to each ancestor due to pedigree collapse. Therefore, for distant relationships, it is more appropriate to conceptualize the relationship inference problem in the form shown in Figure 5B.

The goal of relationship inference under the model in Figure 5B is not to infer “the relationship” between *i* and *j* since there is not just one relationship. Instead, the goal is to infer any one of several quantities of interest such as (1) the most recent genealogical relationship, (2) the most recent genealogical relationship that resulted in detectable shared IBD, (3) the number of genealogical ancestors at each generation in the past, and (4) the number of genealogical ancestors at each generation in the past who contributed observed IBD, or some other suitable quantity that reflects the fact that individuals share many common ancestors through many different relationships.

To derive a relationship estimator for deep relationships, we conceptualize inheritance under the model in Figure 5B and our goal is to infer a statistic that captures this kind of multi-ancestral relationship. Of the statistics above, Statistics 3 and 4 do the most comprehensive job of reflecting the reality of relatedness. However, compared with Statistics 1 and 2, Statistics 3 and 4 pertain to a much larger state space and are therefore more computationally challenging. They are also more susceptible to the statistical problem of non-identifiability or near-nonidentifiability. In particular, many genealogical relationships can give rise to similar observed IBD patterns, so we may be unable to say with high certainty which ancestral relationships gave rise to the observed patterns.

We investigate Statistics 2 and 4 in Section 5.5. There, we show how to infer the number of detectable-IBD-transmitting common ancestors, along with which generation in the past they are from. This is effectively Statistic 4. If we take the minimum generation of such a common ancestor, we get Statistic 3.

Figure 6A shows a comparison of Statistic 3 with the true degree of relationship for individuals who are truly related through only a single common ancestors. Because the estimator of Section 5.5 has a very large state space when the number of observed segments is large, it is computationally taxing for the inference of close relationships that share many IBD segments. Therefore, Figure 6A excludes close relationships. From Figure 6 it can be seen that Statistic 3, the degree induced by the most recent detectable-IBD-transmitting common ancestor, tracks reasonably well with the true degree of relationship, although it is noisy.

**Figure 6.**
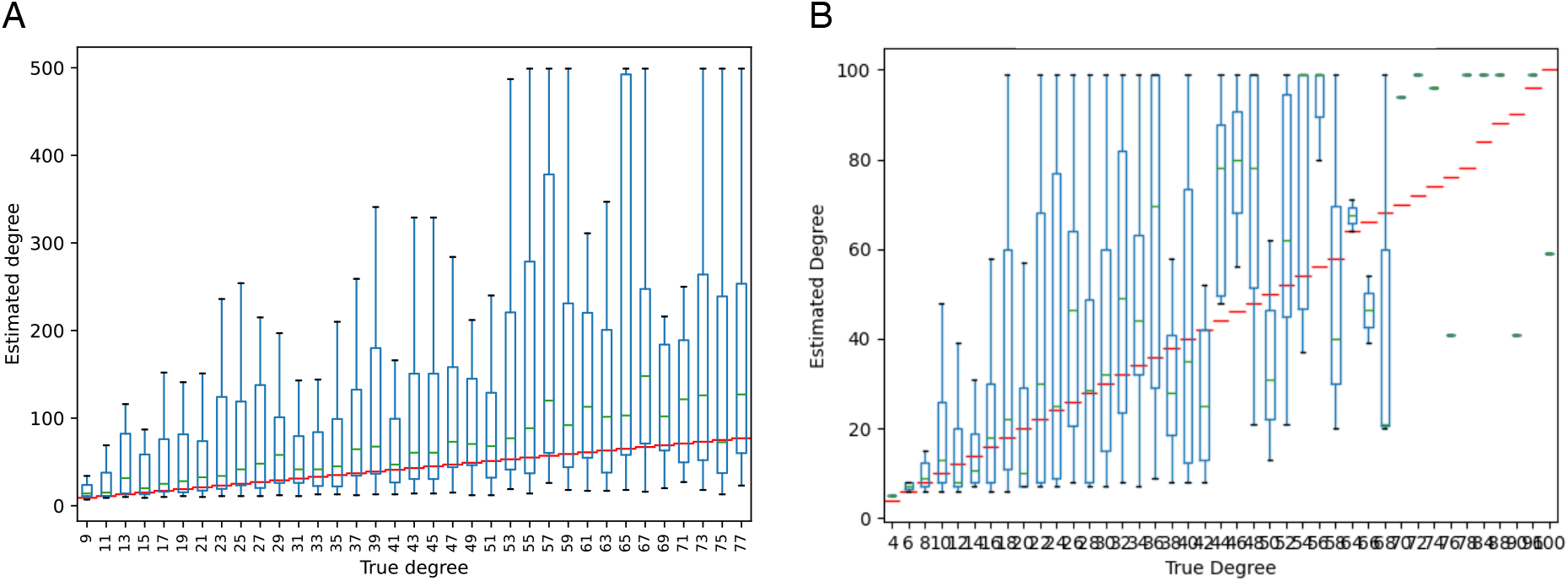
(A) Concordance between Statistic 3 (the degree induced by the most recent detectable-IBD-transmitting common ancestor) and the true degree for relationships in which two people were truly related through a single pair of common ancestors. (B) Concordance between Statistic 3 and the shortest degree among individuals related through multiple common ancestors.

For a more realistic application of the multi-ancestor estimator, we simulated 500 relative pairs in a population with effective size *N* = 5, 000, leading to multiple common ancestors, on average, contributing more than 5 cM. Figure 6B shows a comparison of Statistic 3 (the inferred degree of the most recent detectable-IBD-contributing common ancestor) with the true degree of the most recent detectable-IBD-contributing common ancestor. For these estimates, we restricted the state space to include at most three common ancestors and generation depths of at most 100 generations. Despite these restrictions, Figure 6B shows that the estimator tracks reasonably well with the true degree, although the estimates are quite noisy as expected.

Estimators based on the model in Figure 6B have an important advantage over estimators based on the model in Figure 6A. Specifically, estimators based on Figure 6A, model background IBD segments as noise that confounds the signal of the close relationship (orange curve) on which we are focusing. The conceptualization of background IBD as noise effectively imposes a ceiling on the depth of relationships that can be inferred because a short segment whose length is close to the expected background length cannot be distinguished from background noise. The likelihood is maximized by assigning such segments to the “noise” category, rather than assigning them to distant ancestors. In contrast, estimators like the one shown in Figure 6 treat all segments as real and they do not have such a ceiling.

## 5. Methods

### 5.1 Updating existing relationship estimators by conditioning on observed IBD

Here, we examine how existing relationship estimators (operating under the model in Figure 5A) are affected by conditioning on the event that at least one segment is observed. This allows the direct comparison of existing estimators with versions that condition on observing IBD (Figure 3). We are specifically interested in deriving the distribution of the number and lengths of observed IBD segments, conditional on observing at least one segment.

Following the notation of Ko and Nielsen [2017], let *R* denote a particular relationship between individuals, *i* and *j*, where *R* = (*u, v, a*) indicates that *i* and *j* are related through *a* ∈ {1, 2} common ancestor(s) with *u* meioses separating *i* from the ancestor(s) and *v* meioses separating *j* from the ancestor(s). The total number of meioses is *m* = *u* + *v* and the degree is *d* = *m* − *a* + 1. Let *n* denote the number of segments shared between relatives *i* and *j* arising through relationship *R*. In the conceptual model underlying the relationship inference problem (Figure 5A), some of these segments come from the common ancestor who contributed detectable IBD to *i* and *j*, giving rise to the relationship that we are attempting to infer. Other segments come from other ancestors. Let *n*_*d*_ denote the number of segments that came from the common ancestor of interest and let *n*_*b*_ denote the number of segments that came from other ancestors.

Let {𝓁_1_, …, 𝓁_*n*_} denote the lengths of the *n* = *n*_*d*_ + *n*_*b*_ IBD segments observed between *i* and *j* in units of centimorgans. Let *O* be the event that *i* and *j* are observed to share at least one segment of IBD. Our goal is to compute the probability ℙ (𝓁_1_, …𝓁_*n*_|*O*; *m, a*) of the observed IBD, conditional on the event, *O*, that at least one IBD segment is observed. Assuming that the *n*_*d*_ segments were transmitted through the relationship *R*, this probability is a function of the number, *a*, of most-recent common genealogical ancestors and the number, *m*, of meioses that separate *i* and *j*.

We closely follow the derivation in Huff et al. [2011], who derived the corresponding probability distribution in the unconditional case. As in Huff et al. [2011], we make the simplifying assumption that the *n*_*d*_ segments coming from the most-recent IBD-contributing common ancsestor(s) are the longest segments. This assumption allows us to avoid conditioning on the subset of IBD segments that arose from this ancestor, which allows us to avoid a summation over all subsets of segments that could have come from the common ancestor. Given this simplifying assumption, the distribution of segment lengths is

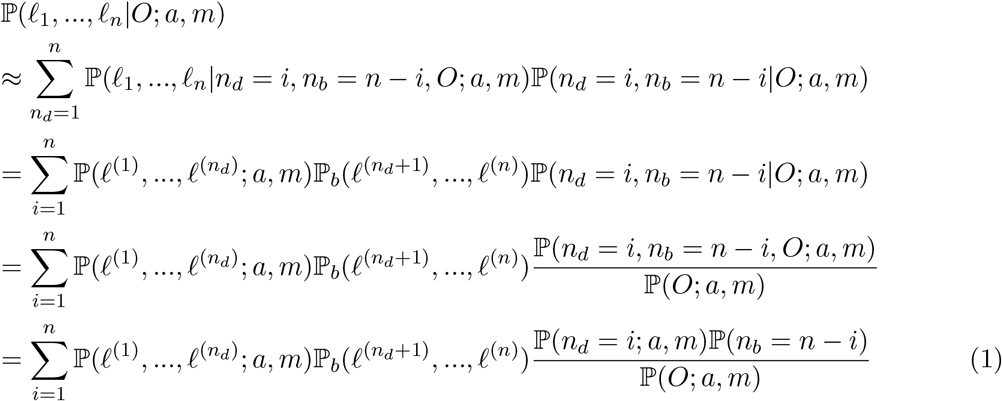

where ℙ_*b*_(·) denotes the probability distribution of background IBD segment lengths.

The terms in Equation (1) can be obtained using equations from Huff et al. [2011] and are as follows:

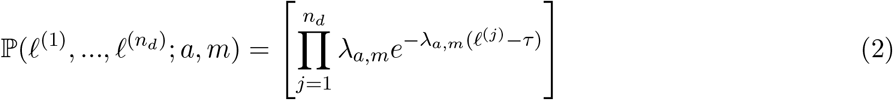

where *λ*_*a,m*_ is the inverse of the expected length of an IBD segment between two people separated by *m* meioses and *τ* is the minimum detectable segment length in centimorgans [Huff et al., 2011]. As in Huff et al. [2011], the segment lengths are modeled as independent, which is likely to be an accurate approximation when *m* is moderate to large [Huff et al., 2011, Caballero et al., 2019]. We are primarily concerned here with distant relationships^1^ so we will make use of the approximation *λ*_*a,m*_ ≈ *m/*100, in which case the right-hand side of Equation (2) doesn’t depend on *a*.

Similarly, the term 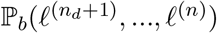 can be modeled as the product of *n* − *n*_*d*_ independent exponential distributions:

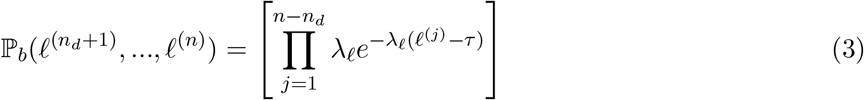

where *λ*_𝓁_ can be found empirically by assuming that most pairs of individuals in a large database are only distantly-related and collecting the lengths of IBD segments shared between many randomly sampled pairs.

The distribution ℙ (*n*_*b*_ = *i*) of the number, *n*_*b*_, of background segments can be found empirically as well. In particular, it is reasonable to model *n*_*b*_ as a Poisson random variable

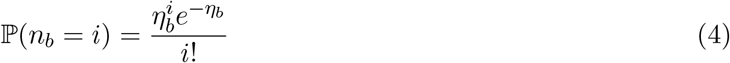

with mean *η*_*b*_ equal to the average number of IBD segments observed between a randomly-sampled pair of individuals in a dataset.

Following Huff et al. [2011], we model the number of observed segments from the genealogically most recent IBD-contributing common ancestor(s) as Poisson, with mean *η*_*a,m,τ*_ :

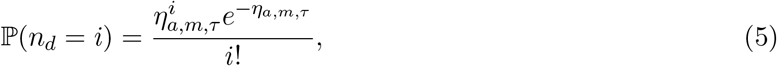

where, for moderate-to-large *m, η*_*a,m,τ*_ can be approximated [Thomas et al., 1994, Huff et al., 2011] as

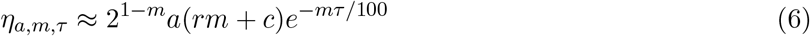

where *r* is the expected number of recombination events per meiosis in the genome, *c* is the number of chromosomes, and *τ* is the minimum detectable segment length. For the autosomal genome in humans, we have *r* ≈ 35 and *c* = 22 [McVean et al., 2004, Huff et al., 2011].

Finally, the probability of observing any segments at all is:

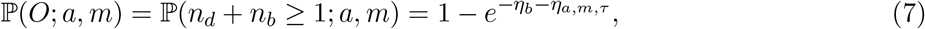

which comes from the fact that *n*_*d*_ and *n*_*b*_ are each Poisson, so their sum is Poisson with mean equal to the sum of the individual means. Equation (7) is one minus the probability that the sum *n*_*d*_ + *n*_*b*_ is zero.

All together, Equation (1) becomes

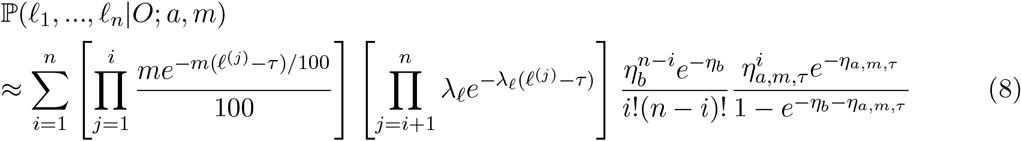

where *λ*_𝓁_ and *η*_*b*_ are found empirically and *η*_*a,m,τ*_ ≈ 2^1−*m*^*a*(*rm* + *c*)*e*^−*mτ/*100^.

Note that Equation (8) is nearly identical to its unconditional version presented in Equation (9) of Huff et al. [2011]. The only difference is the normalizing factor 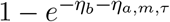 and the fact that the summation starts at 1 rather than at 0. This provides a simpler alternative derivation, as the conditional distribution is effectively obtained from the unconditional expression by ignoring the mass at zero and renormalizing the mass above zero.

### 5.2 The distribution of the total length of IBD

We can also obtain the distribution of the total length of observed IBD. We make the simplifying assumption that there are no background IBD segments in order to obtain a formula that yields values that are comparable the distributions obtained using simulations in existing methods [Henn et al., 2012, Ball et al., 2016]. The joint distribution of the total length of IBD *L* and the number of segments *n* can be obtained by noting that the sum of exponential random variables follows a gamma distribution. Conditioning on the event *O* that at least one segment is observed, we obtain

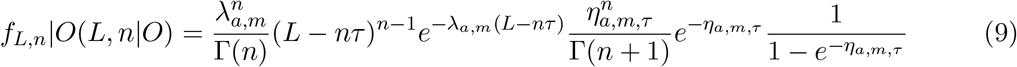

where *η*_*a,m,τ*_ = 2^1−*m*^*a*(*rm* + *c*)*e*^−*mτ/*100^ and *λ*_*a,m*_ ≈ *m/*100.

We can also obtain the distribution of *L* alone by summing over *n*:

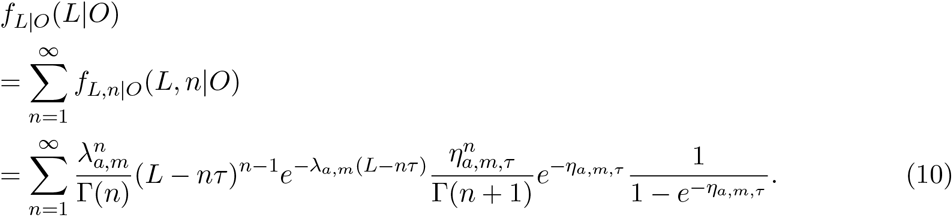

For the case in which the minimum segment length is *τ* = 0, this equation has a closed form, which can be obtained by rearranging the terms to resemble the summation in a modified Bessel function:

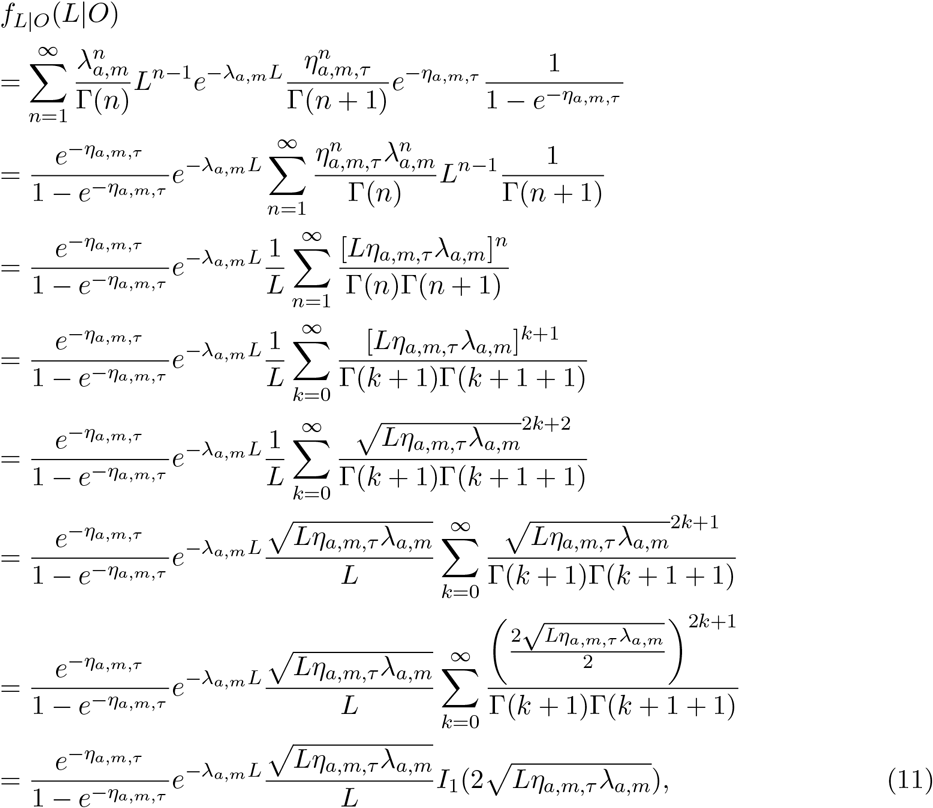

where *I*_1_(·) is the modified Bessel function of the first kind.

Note that the only aspect of Equation (11) that corresponds to conditioning on the event *O* is the factor 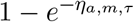, in the denominator. Thus, to obtain the unconditional distribution of the total length, we simply remove this factor:

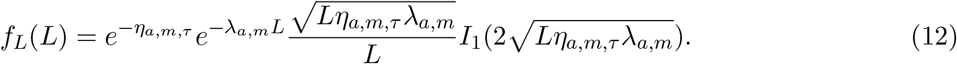

Equation (12) provides the analytical version of the of the empirical distribution that is used by genetic testing companies for relationship inference (for example, see Figure 3 of Henn et al. [2012] and the figure in Section 5.2 of Ball et al. [2016]). Plots of Equations (11) and (12) are shown in Figure 4.

Genetic testing companies typically obtain Equation (12) empirically using simulations. However, because there is a lot of noise in empirical simulations, the distributions obtained by genetic testing companies are noisy. In comparison, the analytical distribution has no noise.

### 5.3. Regions where the likelihood is maximized

From Equation (12), we can analytically obtain the thresholds on *L* that are used for relationship inference by some genetic testing companies. These thresholds are points *L*_*d*_ that partition the space of the total IBD length into regions (*L*_1_, *L*_0_], (*L*_2_, *L*_1_], (*L*_3_, *L*_2_], etc. such that the most likely degree of relationship in the range (*L*_*d*_, *L*_*d*−1_] is *d*. We can also extend these regions to the case of conditional likelihoods using Equation (11).

The range (*L*_*d*_, *L*_*d*−1_] depends only on the degree *d* = *m* − *a* + 1, rather than explicitly on *a* and *m*. For distant relationships, this dependence on *d* alone is reasonable because a relationship of degree *d* with two common ancestors *a* = 2 has a very similar pattern of shared IBD compared with a relationship of degree *d* and one common ancestor *a* = 1. Although age information can help to determine the number of ancestors *a* for close relationships, for very distant relationships, there is almost no available information to determine whether the number of ancestors is 2 or 1. Hence, the degree *d* is often the most relevant quantity to infer for distant relatives.

Suppose for simplicity that *a* is equal to 1. The case *a* = 2 yields similar values of *L*_*d*_ for large *d*. Under this assumption we have *m* = *d* and Equation (12) becomes

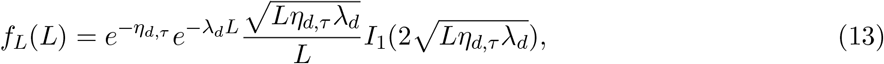

and Equation (11) becomes

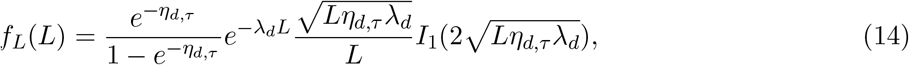

where *λ*_*d*_ = *d/*100 and *η*_*d,τ*_ = 2^1−*d*^(*rd* + *c*)*e*^−*dτ/*100^. Using Equation (13), we can analytically obtain the points *L*_*d*_ by solving for the value of *L* such that *f*_*L*_(*L*; *d* + 1) = *f*_*L*_(*L*; *d*). Using Equation (14) we can obtain these bounds in the conditional case in which at least one segment of IBD is observed. Solving these equations gives the bounds *L*_*d*_ shown in Table (2).

### 5.4. The prior distribution of the generation in which a detectable-IBD-transmitting common ancestor lived

When the minimum observable segment length is *τ* cM, we say that an ancestor is a detectable-IBD-contributing ancestor for a pair of individuals *i* and *j* if they contributed at least one IBD segment longer than *τ* cM to *i* and *j*. To obtain the expected number of detectable-IBD-contributing common ancestors at each generation in the past, we first obtain a more fundamental quantity, which is the expected number *E*[*S*_*g*_] of detectable transmitted IBD segments arising in generation *g*. To obtain *E*[*S*_*g*_], note that all of the DNA in a person living in the present day came from their set of ancestors living in generation *g* in the past. Moreover, *g* generations in the past, each copy of a present-day individual’s genome existed in approximately *H*_*g*_ tiny semi-independent haplotype chunks that would ultimately combine over the generations to produce the linear genome of the present-day person. The expected number *E*[*H*_*g*_] of haplotypes chunks is

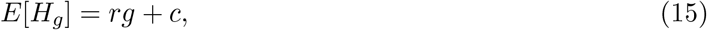

where *c* is the number of autosomes under consideration (e.g., *c* = 22 for humans) and *r* is the expected number of recombination events per meiosis per genome (e.g., *r* ≈ 35 cM for humans). Equation (15) comes from the fact that a genome has, on average, *rg* + *c* − 1 breaks introduced into it over *g* generations including the obligatory breaks at chromosome endpoints, yielding *rg* + *c* − 1 + 1 = *rg* + *c* pieces on average. This result comes from the backward-in-time version of the reasoning of Thomas et al. [1994], who considered the number of segments a chromosome breaks into as it is transmitted forward in time.

Now, if we consider the overlapping chunks of two linear genomes *g* generations in the past, the *rc* + *g* chunks from one copy of *i*’s genome will overlap the *rc* + *g* chunks from a copy of *j*’s genome in an average of 2(*rc* + *g*) overlapping regions. The length one of these overlapping chunks between two people found anywhere in the genome is exponentially distributed with mean 100*/*2*g*. Therefore, the probability that a shared segment anywhere in the genome is longer than *τ* cM is then *e*^−2*gτ/*100^ = *e*^−*gτ/*50^. Finally, under the coalescent model, the probability that a shared segment between two individuals coalesces *g* generations in the past in a population with *N* diploid individuals is 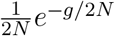. Putting all of this together, we get

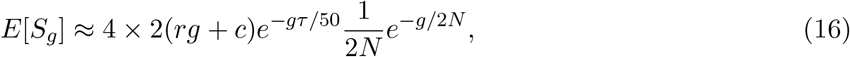

where the factor of 4 comes from the fact that individuals *i* and *j* each have two copies of the linear genome that can coalesce with one another.

We now use Equation (16) to find the expected number of distinct detectable-IBD-transmitting ancestors in generation *g*. If *i* and *j* share *s* segments through common ancestors living *g* generations ago, then the probability they came from exactly *k* distinct common ancestors can be obtained as follows: let *c* be the number of common ancestors. The number of ways of placing *s* indistinguishable segments into *c* individuals is 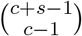 [Ross, 2002]. The number of ways of choosing *k* specific individuals is 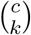. Finally, the number of ways of placing *s* indistinguishable segments into *k* specific individuals so that each individual has at least one segment is 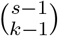 [Ross, 2002]. Note that each of the *s* shared segments must have come from a common ancestor. So, all together, the probability that there are *k* distinct ancestors in generation *g*, given that there are *s* shared segments is

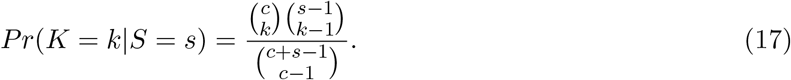

Thus, the expected number of common ancestors, given that there are *s* segments is

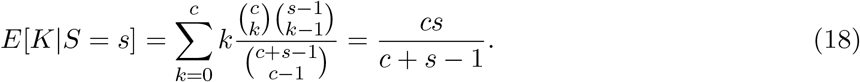

Modeling the number of shared segments *S*_*g*_ arising in generation *g* as Poisson distributed with mean *E*[*S*_*g*_], we find that the expected number of distinct common ancestors *K*_*g*_ is

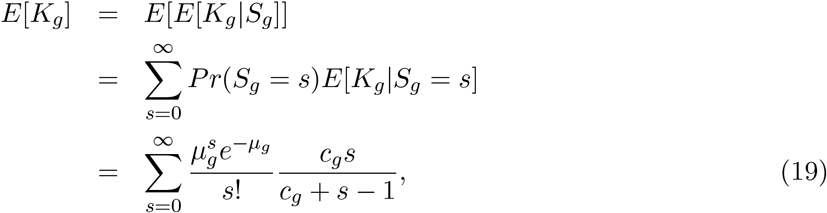

where 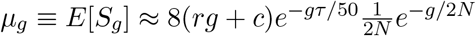.

### 5.5. Multi-ancestor relationship inference

We noted in Section 4 that the existing modeling framework is not particularly appropriate for inferring deep relationships because this framework assumes that people have just one common ancestor, or one mating pair of ancestors who transmitted IBD. However, the prior distributions shown in Figure 2 demonstrate that it is not unusual for a pair of individuals to have multiple detectable-IBD-transmitting ancestors at various generations in the past.

As we noted, several statistics are more appropriate for capturing the complexity of true relationships, such as 1) the time to the most recent genealogical ancestor, 2) the time to the most recent IBD-contributing common ancestor, 3) the number of genealogical ancestors in each generation in the past, and 4) the number of detectable-IBD-transmitting common ancestors at each generation in the past. Statistics 2 and 4 are perhaps easier to infer because they pertain to ancestors who contributed observed IBD. We focus on these statistics to avoid the extra layer of complexity involved in inferring ancestors who left no genetic traces of themselves in the putative relative pair. Statistics 1 and 3 are likely to depend largely on the demographic history of the population and are left for future work.

Suppose we observe *n* segments with a total of *L* cM shared between a pair of individuals. What is our best estimate for the number of ancestors who transmitted these segments and the generations in which they lived? To answer this question, we can write down the probability of the observed IBD, given the full, complex ancestral relationship of the two individuals.

Suppose the two people share a total of *K* common ancestors who lived in generations *g* ∈ 𝒢 in the past for some set of generations 𝒢. Let *k*_*g*_ be the number of ancestors in generation *g*, so that we have Σ_*g*∈*𝒢*_ *k*_*g*_ = *K*. Let *d*_*g*_ be the degree separating a pair of individuals with a common ancestor in generation *g*. For example if the two people are contemporaneous, then *d*_*g*_ = 2*g*, but they need not be contemporaneous.

We can find the joint distribution of *n* and *L* by summing over the number of segments attributable to ancestors in each generation. Let *n*_*g*_ be the number of segments inherited from the ancestors in generation *g*. The generation from which a segment was inherited specifies the expected length *θ*_*g*_ of the segment. Since the segment lengths are exponentially distributed, the expected total length *L*_*g*_ of the *n*_*g*_ segments arising in generation *g* follows a Gamma distribution with shape parameter *n*_*g*_ and scale parameter *θ*_*g*_.

The total observed length *L* = Σ_*g*∈𝒢_ *L*_*g*_ can be approximated as the sum of |*G*| independent but not identically distributed gamma-distributed random variables. *L* can be approximated by a single gamma-distributed random variable whose mean and variance match the mean and variance of the sum Σ_*g*𝒢_ *L*_*g*_ [Covo and Elalouf, 2014]. Since the *L*_*g*_ are independent, the mean and variance of *L* are

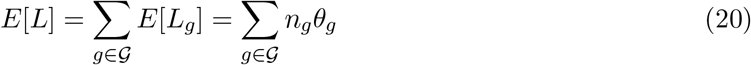

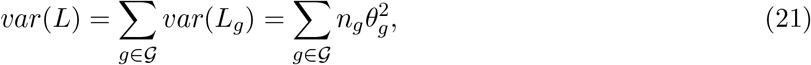

where we have used the fact that the parameters *α* and *β* of a gamma-distributed random variable *X* satisfy *E*[*X*] = *αβ* and *var*(*X*) = *αβ*^2^. It follows that the parameters *α*_*L*_ and *β*_*L*_ of *L*, given {*n*_*g*_}_*g*∈𝒢_ satisfy

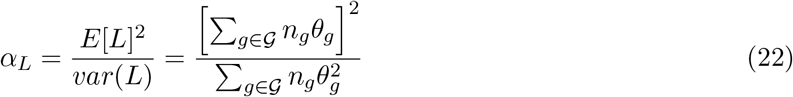

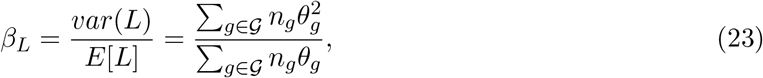

where the mean segment length *θ*_*g*_ from an ancestor in generation *g* is 100*/d*_*g*_ and *d*_*g*_ is the degree of the relationship between the two relatives *i* and *j* that passes through an ancestor in generation *g*. Given that {*n*_*g*_}_*g*∈𝒢_ ≡ **n** segments are transmitted from each generation *g* ∈ 𝒢 and that the degrees of the induced relationships are {*d*_*g*_}_*g*∈𝒢_, we find that the observed number and total length of IBD has the distribution

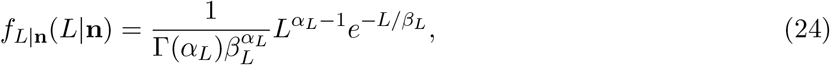

where *α*_*L*_ and *β*_*L*_ are given by Equations (22) and (23).

We now need to know the probability of **n** ≡ {*n*_*g*_}_*g*∈𝒢_ conditional on **k** ≡ {*k*_*g*_}_*g*∈𝒢_ and the fact that each ancestor in **k** contributed at least one segment longer than *τ*. Since there are *E*[*K*_*g*_] ancestors on average in generation *g* and an expected number *E*[*S*_*g*_] of IBD segments, that means that each ancestor contributes *E*[*S*_*g*_]*/E*[*c*_*g*_] segments on average, where *c*_*g*_ is the number of genealogical common ancestors in generation *g*. If there are *k*_*g*_ ancestors in generation *g* then the expected number of contributed segments, beyond the *k*_*g*_ obligatorily contributed segments is *k*_*g*_*E*[*S*_*g*_]*/E*[*c*_*g*_], where *c*_*g*_ is the expected number of distinct common ancestors in generation *g*.

Putting this all together, we find that

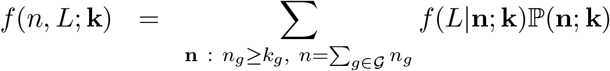

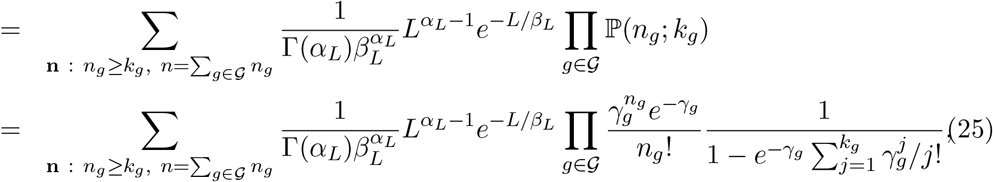

where we have modeled the *n*_*g*_ segments from the *k*_*g*_ ancestors in generation *g* as Poisson distributed with mean *γ*_*g*_, conditional on the event that *n*_*g*_ ≥ *k*_*g*_. Here, *γ*_*g*_ is given by

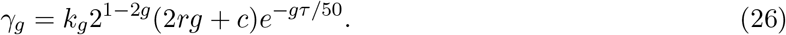

We can also use a value of *γ*_*g*_ that more accurately accounts for the fact that each common ancestor is actually connected to their descendants through multiple lineages. Such a value of *γ*_*g*_ is useful when working with populations with background IBD in which the expected number of segments transmitted from an ancestor in generation *g* can be much higher than in a large population with low amounts of background IBD. In this case, each common ancestor in generation *g* contributes *E*[*S*_*g*_]*/E*[*c*_*g*_] segments, on average, where *c*_*g*_ is the number of common ancestors shared between *i* and *j* in generation *g* in the past. Since there are *k*_*g*_ ancestors in our estimator, this gives

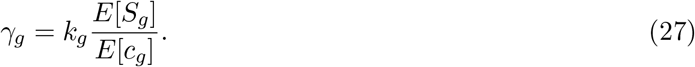

The quantity *c*_*g*_ is derived in Appendix A.

Given counts **k** = {*k*_*g*_}_*g*∈𝒢_ of IBD-contributing ancestors in each generation in the past, Equation (25) gives the probability of observing *n* segments of total length *L*. By viewing Equation (25) as a likelihood, we can infer the most likely set of ancestral relationships that gave rise to the observed IBD and by employing the prior in Equation (19), we can obtain a Bayesian estimate. Since the parameter space of possible relationships is enormous, this approach is likely to be infeasible for relationships sharing more than a few segments. We leave it to future work to make versions or approximations of this estimator that are more efficient.

## 6. Discussion

In this paper, we have shown that existing relationship estimators that do not condition on the event that IBD is observed between a pair of relatives produce biased estimates, inferring all sufficiently distant relationships to be ten degrees. This is a fundamental property of the likelihood of the segment data and it affects all estimators that are explicitly or implicitly based on unconditional distributions of segment lengths or the total IBD. We have also demonstrated that IBD-sharing relationships of degree greater than ten are ubiquitous, amounting to a large percentage of all relationships that are detectable in the population.

Because relationship estimators all demonstrate this bias and because it was supposed that distantly-related individuals are very unlikely to share IBD at all, it has generally been assumed that most relationships are within ten degrees (5^*th*^ cousins) and that relationships beyond seventeen degrees (8^*th*^ cousins) are simply undetectable. The belief that relationships beyond 8^*th*^ cousins were undetectable is evidenced by the fact that relationship estimators used by direct-to-consumer genetic testing companies do not attempt to detect more distant relationships. For example, 23andMe considers IBD sharing down to 20 cM, calling all other relationships as “distant cousins” while the estimators used by AncestryDNA are trained only for 10^*th*^ cousins and closer [Ball et al., 2016]. The fact that existing estimators did not detect very distant relationships was taken as evidence of a fundamental truth about relatedness, rather than raising suspicions about the estimators themselves. The distribution we obtain for the expected number of detectable-IBD-contributing common ancestors at each generation in the past suggests that detectable ancestral relationships sharing a common ancestor 100 or more generations in the past are likely to be common, especially in large genotyping datasets with millions of sampled individuals. The fact that shared segments can be detected from deep relationships is probably not news to the community of researchers working on problems involving deep coalescence. After all, methods such as the PSMC [Li and Durbin, 2011] have leveraged these kinds of deep relationships for years. However, there has been a disconnect between this coalescent-style research and pedigree inference.

The implication of this work is that a large fraction of distant relationship estimates reported by direct-to-consumer genetic testing companies are simply incorrect. Individuals do indeed have many thousands of fifth cousins as these platforms report. However, a large proportion of relatives that are reported as fifth cousins are in fact much more distant.

The reality is perhaps more interesting than the current incorrect estimates imply: we can detect relationships with common ancestors who lived before major world migrations such as European contact in the Americas and the Transatlantic slave trade and before events such as the rise of the Roman Empire and perhaps even the building of the Pyramids at Giza. This finding is particularly important for projects connecting present day people of African ancestry in the United states with relatives living in Africa with whom they share a common ancestor prior to or during the Transatlantic Slave Trade [David, 2023, 2024]. These studies provide a means of uncovering a genealogical history that was lost due to slavery.

Although the variance in distant estimates is high, estimates can be used in aggregate to obtain a more accurate picture of the ancestral connections of an individual, as well as the interrelationships among populations over a timespan of several thousand years. The tapestry of relationships that we can infer may in fact be quite rich.

## 7. Acknowledgements

I would like to thank the employees and research participants of 23andMe who made this research possible. I am especially grateful to Amy L. Williams, William A. Freyman and David A. Hinds for their insightful comments and thoughtful reviews of this manuscript and Peter R. Wilton for insightful points raised in discussions. Funding for this work was provided by NIH grant R35 GM133805 and by 23andMe, Inc. Members of the 23andMe Research Team are Stella Aslibekyan, Adam Auton, Elizabeth Babalola, Robert K. Bell, Jessica Bielenberg, Ninad S. Chaudhary, Zayn Cochinwala, Sayantan Das, Emily DelloRusso, Payam Dibaeinia, Sarah L. Elson, Nicholas Eriksson, Chris Eijsbouts, Teresa Filshtein, Pierre Fontanillas, Davide Foletti, Will Freyman, Zach Fuller, Julie M. Granka, Chris German, Éadaoin Harney, Alejandro Hernandez, Barry Hicks, David A. Hinds, M. Reza Jabalameli, Ethan M. Jewett, Yunxuan Jiang, Sotiris Karagounis, Lucy Kaufmann, Matt Kmiecik, Katelyn Kukar, Alan Kwong, Keng-Han Lin, Yanyu Liang, Bianca A. Llamas, Aly Khan, Steven J. Micheletti, Matthew H. McIntyre, Meghan E. Moreno, Priyanka Nandakumar, Dominique T. Nguyen, Jared O’Connell, Steve Pitts, G. David Poznik, Alexandra Reynoso, Shubham Saini, Morgan Schumacher, Leah Selcer, Anjali J. Shastri, Jingchunzi Shi, Suyash Shringarpure, Keaton Stagaman, Teague Sterling, Qiaojuan Jane Su, Joyce Y. Tung, Susana A. Tat, Vinh Tran, Xin Wang, Wei Wang, Catherine H. Weldon, and Peter Wilton.

## Appendix A. The expected number of distinct common ancestors

Here, we derive the distribution of the number of distinct common ancestors that two present-day people share in generation *g* in the past. Let *a*_*g*_ denote the number of *distinct* ancestors in generation *g*. Assuming a randomly-mating population in which the two parents of each individual in generation *g* − 1 are chosen uniformly at random from among the *N* individuals in the population in generation *g*, we model *a*_*g*_ as the number of distinct draws from a population of *N* distinct individuals when 2*a*_*g*−1_ samples are taken with replacement. We now develop a recurrence relation that approximates the expected number of distinct common ancestors shared between two people in generation *g* in the past.

Let *δ*_*g*_ denote the number of distinct ancestors of individual *i* in generation *g* that are not common ancestors of individual *j*. Let *c*_*g*_ denote the number of distinct common ancestors shared between individuals *i* and *j* in generation *g*. Finally, let 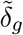 denote the number of distinct ancestors of the *δ*_*g*−1_ individual ancestors of *i* in generation *g* − 1 before considering mergers with ancestors of *j* and let 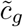 denote the number of distinct ancestors of the *c*_*g*−1_ common ancestors before considering mergers among individuals. Thus 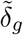is the number of distinct objects when drawing 2*δ*_*g*−1_ samples with replacement from a population of *N* individuals and 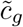 is the number of distinct individuals selected when drawing 2*c*_*g*−1_ objects with replacement from a population of *N* individuals.

The quantity 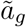 can be obtained from the quantity *a*_*g*−1_ by noting that the probability that any given one of the *N* individuals in generation *g* is a parent of one of the *a*_*g*−1_ individuals in generation *g* − 1 with probability 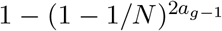. This probability comes from the fact that the individual is selected in a single draw from the population with probability 1*/N*. Thus the probability that they are not selected in any of the 2*a*_*g*−1_ draws is 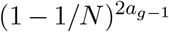. It follows that 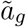 is binary with probability 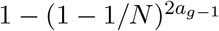 and expectation 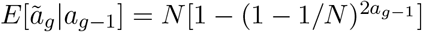. Similarly, 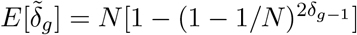 and 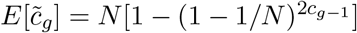.

When the population size is large, the quantities 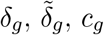 and 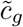 behave quasi-deterministically and we will assume that they follow their expected values. Thus, in generation *g*, the 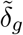 distinct non-common ancestors of *i* overlap with one of the 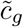 distinct common ancestors at rate 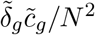 and they overlap with one of the 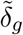 distinct ancestors of individual *j* at rate 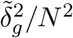. Therefore, the expected number of distinct and common ancestors at generation *g* is

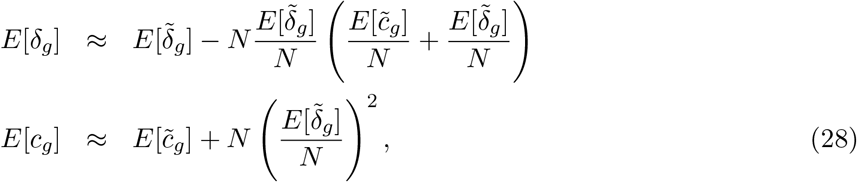

where

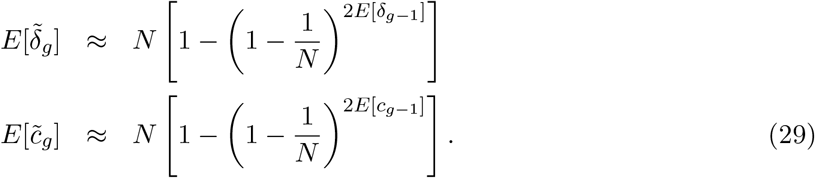

Repeatedly applying this recursion gives an approximation of the number of distinct common ancestors at each generation in the past.

## Footnotes

1 The approximation begins to break down when *a* = 2 and *m* ≤ 3 (avuncular relationships and closer), but in this region of the parameter space, the information coming from the shared IBD is typically so strong that relationships can be inferred accurately even when the likelihood is misspecified.

