## Supplemental Figures for "Correcting model misspecification in relationship estimates"

### S1. Supplemental figures

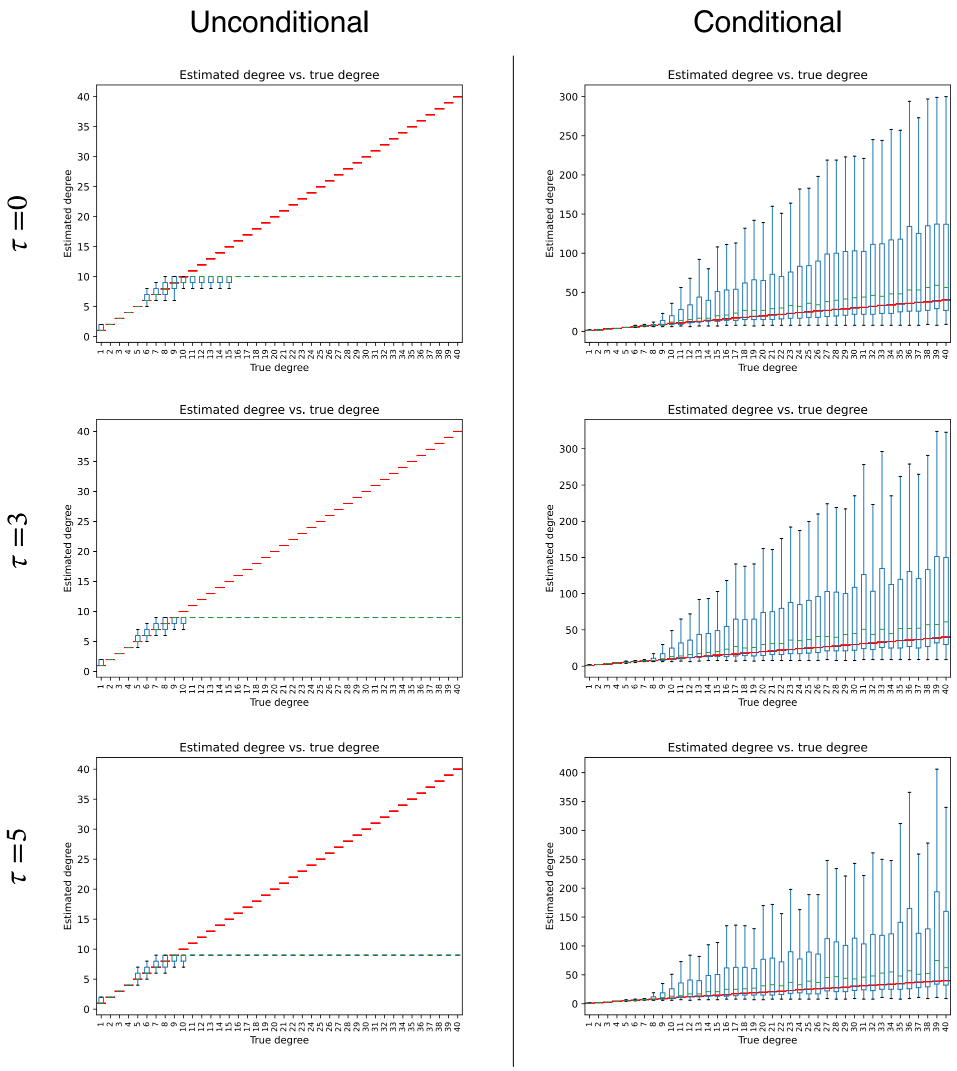


Figure S1. The effect of a minimum segment length $\tau$ on the unconditional and conditional estimators. Including a minimum segment length reduces the number of relationships that share IBD and which can therefore be inferred. However, given that two individuals share IBD segments that exceed the minimum segment length, the accuracy of the estimator is similar to the case in which there is no minimum segment length. This property of the estimators is due to the fact that the exponential distribution of the segment length is memoryless. In other words, given that a segment is longer than $\tau$ its length is distributed according to the same distribution as a segment with minimum length zero. Therefore, the segment contains as much information about the degree of relationship as the un-thresholded segments.
